# Signatures of divergent antimalarial treatment responses in peripheral blood from infants and adults in Malawi

**DOI:** 10.1101/564757

**Authors:** Paul L. Maurizio, Hubaida Fuseini, Gerald Tegha, Mina Hosseinipour, Kristina De Paris

## Abstract

**Background:** Heterogeneity in the immune response to parasite infection is mediated in part by differences in host genetics, sex, and age group. In neonates and infants, ongoing immunological maturation often results in increased susceptibility to infection and variable responses to drug treatment, increasing the risk of complications. Even though significant age-specific effects on host cytokine responses to *Plasmodium falciparum* infection have been identified, age effects on uncomplicated malaria infection and antimalarial treatment remain poorly understood.

**Methods:** In samples of whole blood from a cohort of naturally infected malaria-positive individuals in Malawi (n=63 total; 34 infants <2 years old, 29 adults >18 years old), we assessed blood cytokine levels and characterized monocyte and dendritic cell frequencies at two timepoints: acute infection, and four weeks post antimalarial treatment. We modeled the effects of age group, sex, and timepoint, and evaluated the role of these factors on infection and treatment outcomes.

**Results:** Regardless of treatment timepoint, in our population age was significantly associated with overall blood hemoglobin, which was higher in adults, and plasma nitric oxide, IL-10, and TNF-*α* levels, which were higher in infants. We found a significant effect of age on the hemoglobin treatment response, whereby after treatment, levels increased in infants and decreased in adults. Furthermore, we observed significant age-specific effects on treatment response for overall parasite load, IFN-*γ* and IL-12(p40), and these effects were sex-dependent. We uncovered significant age effects on the overall levels and treatment response of myeloid dendritic cell frequencies. In addition, within each age group, we found continuous age effects on gametocyte levels *(Pfs16*), TNF-*α*, and nitric oxide.

**Conclusions:** In a clinical study of infants and adults experiencing natural malaria infection and receiving antimalarial treatment, we identified age-specific signatures of infection and treatment responses in peripheral blood. We describe host markers that may indicate, and potentially mediate, differential post-treatment outcomes for malaria in infants versus adults.

## Background

Variation in the host response to parasite infection depends on a variety of factors including age, sex, host genetics, pathogen strain, and environment. Infant-associated increases in malaria severity are determined in part by the particularities of the infant immune milieu, making this an important and active area of research [1]. However, in addition to age-dependent effects on infection, effects on the response to anti-parasite chemotherapy are not well understood, even though these effects may impede the global agenda for malaria elimination and eradication [2]. Therefore, our lack of knowledge about age-related differences in immune responses to *Plasmodium falciparum* infection and treatment constrains our ability to develop protective antimalarial vaccines and therapeutics for younger individuals who may be at increased risk for severe complications [3, 4, 5].

In malaria endemic regions, repeated exposure to parasites may generate adaptive immunity in some infant populations as a mechanism for protection from severe disease, after the protection offered by maternal antibodies has waned [6, 7, 8, 9, 10, 11. However, age-dependent changes in immune function may also contribute to improved immune responses in adults. Thus, recent studies have explored age-dependent effects in order to understand the relative contribution of parasitological and host immunological effects on heterogeneity in the response to malaria infection.

Age-dependent effects on the production of anti*-Plasmodium* antibodies against pre-erythrocytic and asexual blood stage antigens were recently reported by Ouédraogo et al. [12]. In addition, in children from Mozambique, significant associations were found between infant age and levels of IgG directed against merozoite-stage *Plasmodium* [13]. Furthermore, age-dependent effects on B cell response magnitude [14] and post-treatment parasite clearance [15] have also been described. Whereas these studies focused on identifying age-dependent differences in adaptive and antibody-related responses to parasite infection, our study focuses on age-specific differences in plasma cytokine levels and monocyte activation-factors that may be critical for determining treatment efficacy in infant populations.

Infants face multiple barriers to overcoming malaria infection, including suboptimal innate immune responses to natural infection and poor antimalarial treatment efficacy, which in some cases results in serious outcomes, such as severe malarial anemia (SMA) or cerebral malaria (CM). Studies have shown that SMA and CM are driven by proinflammatory cytokine secretion and immunopathology, suggesting immunomodulation as a potential avenue for adjunctive therapy to prevent severe outcomes in infants [16, 17, 18, 19]. Although SMA and CM have been a major focus of research in infants, we were interested in identifying age-specific markers of treatment response in *uncomplicated* malaria (UM)—an area that is arguably less well studied and yet remains critical to understanding phenotypic variation in the majority of malaria-infected and treated infants. Therefore, in order to isolate age-specific effects on UM, and also to avoid exacerbation of disease among participants, we excluded individuals who showed evidence of severe anemia from our cohort.

In this study, we examined infant and adult peripheral blood, collected during acute malaria infection and ~4 weeks post-antimalarial treatment, to identify signatures of differential host responses to infection and treatment. Among our main findings, we report significantly higher plasma IL-10 and TNF-*α* levels, and nitric oxide, in infants compared with adults, regardless of treatment. We also observed that IFN-*γ* and IL-12(p40) treatment responses differed significantly based on age, in a sex-specific manner. In addition, we observed several subjects (5 of 63) with apparent treatment failure, or reinfection. Thus, this work improves our understanding of the infant-specific response to malaria infection, implicating inflammatory differences in whole blood treatment responses on post-treatment infection resolution, and may contribute to the development of improved vaccines and therapies for pediatric populations.

## Methods

### Study population and sample collection

We randomly selected subjects for this study from patients who tested positive for *Plasmodium falciparum* infection, February 1st, 2012 through May 22nd, 2012, at the Kamuzu Central Hospital (KCH) outpatient clinic in Malawi. A total of 34 infants (4-24 months) and 29 adults (19-70 years) were enrolled (Table 1). We obtained informed written consent from adult participants and from parents of infant participants during the first clinic visit. Enrollment in the study was voluntary and all infected patients received antimalarial treatment independent of enrollment. The study was approved by the Institutional Review Board at UNC and the National Health Sciences Research Committee, under the oversight of the Ministry of Health, in Malawi. The institutional guidelines strictly adhere to the World Medical Association’s Declaration of Helsinki.

**Table 1:** Clinical characteristics of study participants. Where applicable, values are given as mean (1 s.d.).

|  | Infants |  | Adults |  |
| --- | --- | --- | --- | --- |
| **Acute** |  |  |  |  |
| No. of Participants | 34 |  | 29 |  |
| Age, yrs | 0 – 2 |  | 19 – 70 |  |
| Hemoglobin, g/dL blood | 9.16 | (2.06) | 14.05 | (2.53) |
| Parasite load, $\mu\text{l}^{-1}$ blood ($\times 10^4$) | 7.08 | (12.4) | 1.10 | (2.32) |
| Gametocytemia, $\mu\text{l}^{-1}$ blood | | | | |
| Pfs16 | 36.97 | (47.25) | 44.19 | (40.69) |
| Pfs25 | 2.74 | (8.19) | 0.517 | (1.22) |
| Pfs230 ($\times 10^5$) | 2.24 | (9.83) | 15.6 | (70.2) |
| % positive for anti-*Plasmodium* Ab | 50.00 |  | 79.31 |  |
| **Post-treatment** |  |  |  |  |
| Hemoglobin, g/dL blood | 10.76 | (2.02) | 13.03 | (2.23) |
| Parasite load, $\mu\text{l}^{-1}$ blood ($\times 10^4$) | 0.2589 | (1.338) | 0.0002 | (0.0011) |
| % positive for anti-*Plasmodium* Ab | 44.12 |  | 65.52 |  |

Individuals who visited the hospital and whose clinical diagnosis was consistent with malaria were subsequently screened by a rapid diagnostic test (RDT) to determine malaria positivity, and then enrolled in the study (n=63). Study participants were asked to donate a venous blood sample (infants: 3-5 mL; adults: 10 mL) at their first visit (V1; “acute pre-treatment”). Malaria infection was confirmed by microscopic examination of blood smears. Infants with severe malaria (hemoglobin < 8.0 g/dL and hematocrit < 18%) were excluded from the study to avoid the risk of exacerbating SMA. In addition, whole blood from participants was blotted and dried on Whatman 903™ protein saver cards (#10534612) for gametocytemia analysis.

Infected participants were prescribed antimalarial chemotherapy, which consisted of a first-line regimen of artemether-lumefantrine (AL), and were asked to return in 4-6 weeks for a second visit (V2; “post-treatment”) and blood sample collection. Subjects’ samples and clinical details were de-identified in Malawi. Age, sex, and parasitemia of each patient were recorded with a corresponding unique patient ID code. Blood plasma was collected and stored at -80°C. Peripheral blood mononuclear cells (PBMCs) were isolated using Ficoll-Paque gradient separation and then frozen in 10% DMSO / 90% fetal bovine serum (FBS) and stored in liquid nitrogen. De-identified samples, including blood plasma, PBMCs and dried blood spots, were shipped to the University of North Carolina at Chapel Hill for additional analysis. Details about the selection and phenotyping of study participants are summarized in (Figure 1).

**Figure 1:**
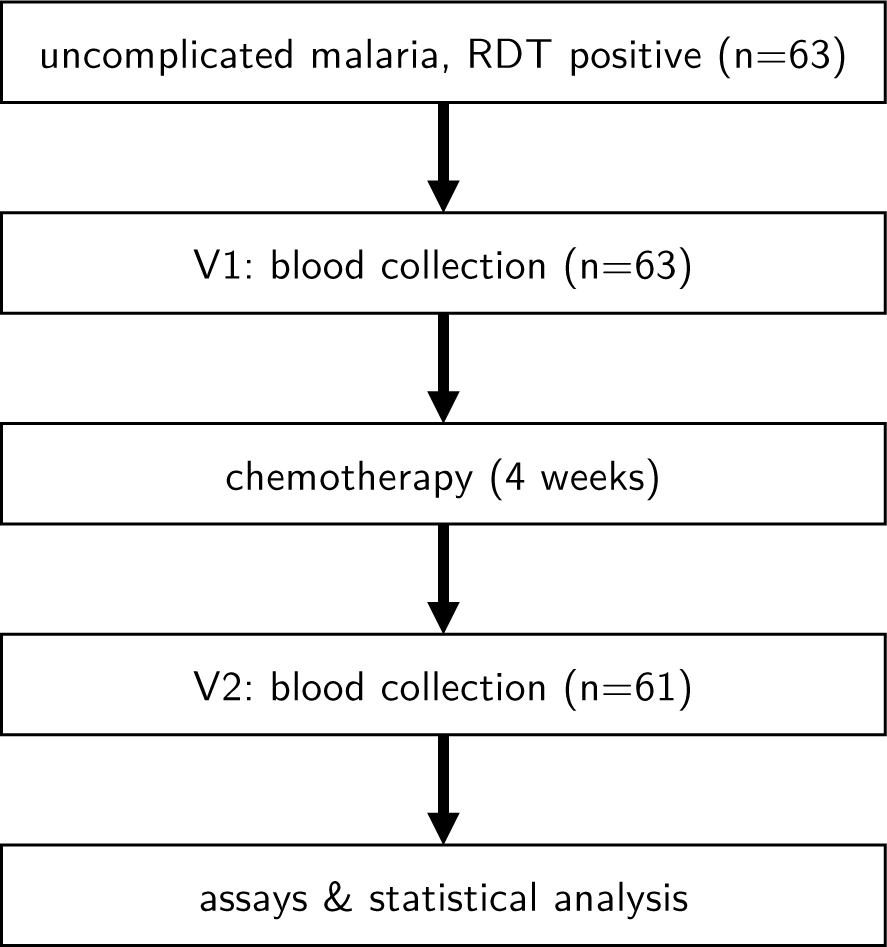
Study population and data collection.

### Parasite load

To determine the level of infection in all malaria-positive subjects, parasitemia was quantified at the Kamuzu Central Hospital clinic in Malawi by light microscopy of thick blood smears at V1 and V2. All slides were read by two expert readers independently and mean values are used as phenotypes; in cases with data discordance, a third reader was assigned.

### Stage-specific gametocytemia

To detect mature parasite infection based on estimated gametocyte load, we used quantitative real-time PCR (qRT-PCR). Total DNA and total RNA were isolated from subjects’ dried blood spots at V1 using nucleic acid purification kits (Norgen Biotek #35300, #36000), and cDNAs were used in qRT-PCR assays [20, 21]. These cDNAs were used to measure RNA expression levels of *P. falciparum* gametocyte-specific genes, including *Pfs25* (mature gametocyte), *Pfs16* (early gametocyte) [22], and *Pfs230*, which encodes a gametocyte antigen important for eliciting host antibodies that inhibit parasite transmission to a mosquito host [23, 24].

### Antimalarial antibodies

We assessed antimalarial antibodies using a semi-quantitative human malaria antibody ELISA kit (IBL International Inc., Hamburg, Germany #RE58901), according to manufacturer’s protocol. From these results, the fraction of infant and adult participants who tested positive for malaria-specific antibodies (IgM or IgG) were calculated.

### Hemoglobin

Hemoglobin levels were measured in clinic, at V1 and V2, and are reported in g/dL.

### Nitric oxide

Plasma samples were deproteinated and NO levels were quantified for V1 and V2 using the QuantiChrom™ nitric oxide assay kit (BioAssay Systems #D2NO-100). Quantification using OD was carried out according to the manufacturer’s protocol (PerkinElmer). Concentrations were based on absorbances normalized to the manufacturer’s standard and calculated via the Beer-Lambert law.

### Plasma cytokines

The following analytes were measured in the plasma, for V1 and V2, using the MILLIPLEX MAP Human Cytokine/Chemokine Magnetic Bead Panel/Immunology Multiplex Assay (EMD Millipore #HCYTOMAG-60K): GM-CSF, IFN-*_γ_* IL-10, IL-12(p40), IL-12(p70), sCD40L, IL-1,*β*, IL-6, and TNF-*α*. Assays were performed according to the manufacturer protocols on a MagPix (Luminex) instrument at the UNC-Chapel Hill Center for AIDS Research (CFAR) HIV/STD Laboratory Core. Standard curves were fit and experimental concentrations determined from a 5-parameter weighted logistic model using the xPONENT_®_ software (v4.1.308.0).

### Monocyte and dendritic cell composition

Flow-cytometric analysis was performed to characterize myeloid dendritic cell (mDC) and monocyte (Mo) frequencies in PBMCs. All antibodies were purchased from BD Biosciences (San Jose, CA). Cells were stained according to BD protocols using the following mouse anti-human antibodies: CD3 (clone SP34-2), CD14 (clone M5E2), CD16 (clone 3G8), CD20 (clone 2H7), CD33 (clone P67.6), HLA-DR (clone G46.6), and CD11c (clone S-HCL-3). MDC frequencies were reported as percentage of mononuclear cells (MNC). Monocytes were further defined by gating as traditional monocytes (CD14^++^CD16^—^), inflammatory monocytes (CD14^++^CD16^+^) and patrolling monocytes (CD14^dim^ CD16^++^) (see Supplemental Figure S1). Samples were acquired on the LSR11 (BD; San Jose, CA) using FACS DIVA software and analyzed with FlowJo (TreeStar, Inc., Ashland, OR).

### Statistical methods

We analyzed our data in the statistical programming language R [25]. Responses were measured for each study participant, using peripheral blood samples collected at two time points: immediately after malaria diagnosis, at visit 1 (V1); and approximately four weeks after completing antimalarial treatment, at visit 2 (V2). Some phenotypes were only measured at V1, and some were measured at both V1 and V2.

We used a zero-inflated Poisson (ZIP) regression model [26] (log link) to evaluate the effect of age and visit on microscopy-based parasite counts at V1 and V2. In brief, ZIP regression uses a two-component mixture model that simultaneously accounts for zero-and non-zero counts using a Poisson, as well as accounting for zero inflation using the binomial distribution (probit link), which is fit using maximum likelihood estimation via the R package pscl [27, 28].

To model effects of age on gametocytemia, as measured from dried blood spots collected during V1 only, we used the exact Wilcoxon-Mann-Whitney two-sample rank-sum test via the R package coin [29], stratifying by sex. We report a two-sided *p*-value.

To model effects of age and sex on antimalarial antibody results (“negative”, “grey”, or “positive”) at V1 and V2, we used ordered logistic regression (a cumulative link model [30]), via the R package MASS [31].

For all additional blood analyte phenotypes that were measured at both V1 and V2, we modeled our data using a rank-based nonparametric model that accommodates longitudinal data which is collected in a factorial design [32, 33]. The model is implemented in the `R` package `nparLD` [34]; ranks were contrasted between groups and used to calculate ANOVA-type statistics [35] according to the factors of interest, which were in our case: age group (infant, adult), sex (male, female), visit (V1, V2), and their pairwise and three-way interactions. Among our subjects, there were missing data points in at least one phenotype: for one individual on the first visit (V1), and for six individuals on the second visit (V2).

## Results

### Subjects

Our study population was comprised of 63 enrolled subjects, including 34 infants < 2 years old (n_females_=16, n_males_=18, and 29 adults ≥ 18 years old (n_females_=16, n_males_=13). All enrolled subjects tested positive for malaria by RDT. Characteristics of the infant and adult participants are provided in Table 1.

### Parasite load

To determine the effect of antimalarial treatment on parasite burden in infected adults and infants, and to test for the effect of age and sex, we quantified parasite loads at V1 and V2 using microscopy of patients’ thick blood smears. During acute infection (V1), parasite loads were detected in 21 of the 27 adults measured (77.8%) and 25 of the 33 infants measured (75.8%), indicating increased sensitivity of RDT-based diagnosis vs. microscopy. Among infants and adults with detectable parasite loads at V1, parasite counts were significantly higher on average (*p <* 10^−16^), by more than 6-fold, in infants (9.35 × 10^4^ μl^−1^) compared with adults (1.40 × 10^4^ μl^−1^); in addition, we found a significant overall effect of age, and a significant age-by-sex interaction (both *p* < 2 × 10^−16^). We found a significant overall zero-inflation intercept (*p* = 0.0225), indicating the detection of excess zero (undetectable) counts in our data set, and these were unaffected by age or sex.

After antimalarial treatment (V2), parasite counts decreased to undetectable in all but 5 female subjects who had residual detectable parasitemia (1 adult, 4 infants). For 4 of these 5, parasite loads nevertheless decreased substantially from V1 to V2 (Figure 2A)

**Figure 2:**
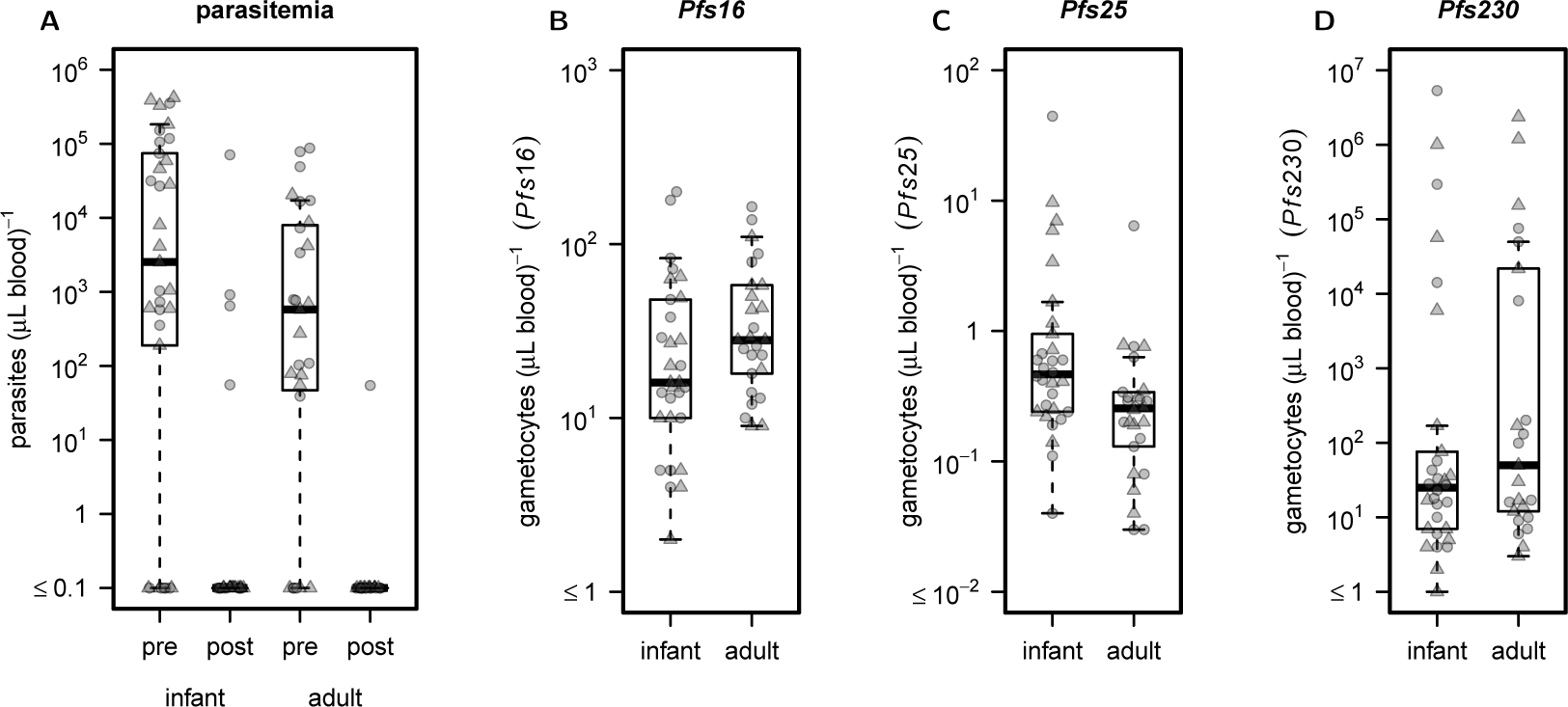
Infant age is associated with increases in blood-stage parasitemia and gametocytemia during acute infection, and incomplete parasite clearance post-treatment. Parasite load (parasites/uL in whole blood) (A) was measured in infants and adults by whole blood microscopy. Gametocytemia was measured using quantitative real-time PCR on cDNAs prepared from dried blood spot RNA, and gametocyte quantification was based on stage-specific expression of the following *Plasmodium falciparum* genes during acute infection: *Pfs16* (early gametocyte) (B); *Pfs25* (mature gametocyte) (C); and *Pfs230* (candidate gametocyte gene for transmission-blocking vaccine) (D), presented as gametocytes/*μ*L. Circular data points indicate female subjects, and triangles indicate male subjects.

### Stage-specific gametocytemia

In the human host, *Plasmodium falciparum* traverses through several life stages. During bloodfeeding at the dermis of a naive host, an infected female *Anopheles* mosquito transmits parasites to the host bloodstream as sporozoites. They mature into schizonts in the liver, rupture from liver cells as merozoites, and grow as ring-stage trophozoites which then enter a schizont-merozoite-trophozoite cycle [36, 37]. A proportion of blood-stage parasites develop into gametocytes, which are the haploid, sexual stage parasites that can subsequently be transmitted to a new female mosquito during bloodfeeding. Whereas it is blood stage parasites that primarily drive clinical disease, gameotocytes are important for human-to-mosquito transmission.

In order to determine the effect of age on stage-specific gametocyte levels in patients’ blood at V1, we estimated *P. falciparum* gametocyte levels in subjects’ blood using qRT-PCR. We used gametocyte stage-specific primers to quantify gene expression of *Pfs16, Pfs25* and *Pfs230* from cDNAs prepared from dried blood spots taken during V1. These quantities are not significantly correlated (*p >* 0.08), indicating that they are likely marking different gametocyte populations in our data set. Whereas, using the *Pfs16* (Figure 2B) and *Pfs230* (Figure 2C) primers, the estimated number of gametocytes did not differ significantly between age groups, we found that infants had significantly higher levels of *Pfs25*-expressing gametocytes (median 0.465 gametocytes/μl) compared with adults (0.255/μl) (*p* = 0.00685) (Figure 2D). We observed no sex-based differences in gene expression for *Pfs16*, *Pfs25* or *Pfs230.*

### Hemoglobin

*Plasmodium* parasites infect erythrocytes, which are a source of hemoglobin (Hb), a host product that is toxic to the parasites. Thus, to survive, *Plasmodium* converts Hb into hemozoin, a chemical more favorable to the parasite. Effects of parasites on Hb levels in the blood may indicate differences in the physiology and composition of host erythrocytes. In order to determine the effect of sex, age, and antimalarial treatment on Hb in study participants, we measured Hb at V1 and V2. We found a significant overall effect of age on Hb levels (higher in adults, *p* = 3.86 × 10^−15^), a significant main effect of sex (higher in females, *p* = 5.6 × 10^−3^), as well as a significant age:visit interaction effect (*p* = 3.14 × 10^−4^) (Figure 3). Compared with the treatment response in adults, whose Hb levels were lower on V2 compared with V1, Hb levels in infants were higher on V2 compared with V1.

**Figure 3:**
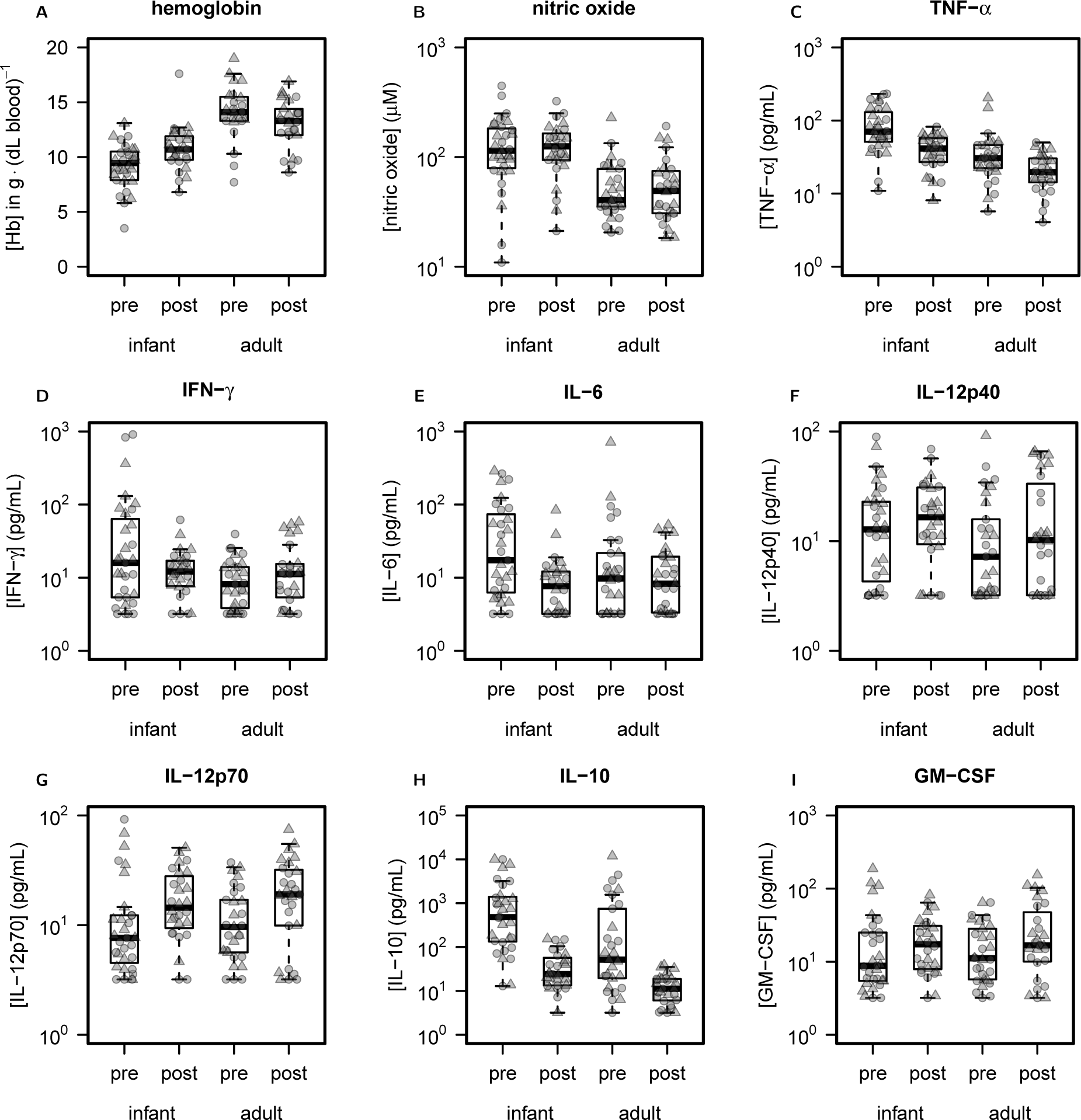
Blood markers in infants differ significantly from adults during acute infection, and respond differentially to antimalarial treatment. The concentrations of the following analytes were assayed, for adult and infant samples collected during acute infection and post-treatment (in pg/mL): (A) TNF-*α*, (B) IFN-*_γ_*, (C) IL-6, (D) IL-12(p40), (E) IL-12(p70), (F) IL-10, and (G) GM-CSF. Levels of (H) hemoglobin in whole blood (g/dL) and (I) nitric oxide (*μ*M) in plasma were also analyzed. Concentrations are presented for acute infection and post-treatment, and stratified by age group. Circles indicate females, and triangles indicate males.

### Antimalarial antibody response

During V1, half of all infants in our study (17 of 34 total; or 10 of 18 males and 7 of 16 females) had detectable antimalarial antibodies, indicating prior exposure to malaria parasites or acquisition of maternal antimalarial antibodies. This is in contrast with the 22 of 29 adults (75.9%; or 10 of 13 males and 12 of 16 females) who had detectable antimalarial antibody at V1, suggesting increased parasite exposure or increased capacity for antibody production, resulting in increased antibody detectability, in adults compared with infants. Thus, we found a significant overall effect of age in our samples (p=0.0298), but no significant effects of sex or treatment. We found that the detectability of antimalarial antibodies was reduced to undetectable levels in five individuals between V1 and V2. Among these five individuals, two were adults (1 male, 1 female) and three were infants (2 males, 1 female). Only two subjects, both infants (1 male, 1 female), transitioned from no detectable antimalarial antibody at V1 to detectable antibody at V2 (Supplemental Figure S2, Supplemental Table S1).

### Nitric oxide

Nitric oxide (NO) is a molecular effector that is released by activated immune cells in their defense against parasite infection [38]. Increased plasma NO levels in adults and children have been associated with protection from malaria [39, 40, 41]. In order to determine whether age, sex, or treatment significantly affected NO levels in our population, we measured NO levels in plasma. We detected significant age-dependent effects on NO levels (*p* = 1.191 × 10^−10^). However, we did not detect significant overall effects of treatment on NO concentrations. No significant sex-specific effect on NO was observed, although the variation in NO at both time points was substantially higher in infant females (*sd*_V1_ = 121.159, *sd*_V2_ = 82.213) than in infant males (*sd*_V1_ = 47.508, *sd*_V2_ = 49.970) (Figure 3).

### Plasma cytokines

To characterize the host immunological response to malaria infection and antimalarial treatment, we measured cytokine protein levels using a MILLIPLEX panel of nine analytes (TNF-*α*, IFN-*_γ_*, IL-6, IL-12 (p40), IL-12 (p70), IL-10, GM-CSF, sCD40L, and IL-1,*β*). To model our data, we used a nonparametric, rank-based statistical framework developed for paired longitudinal measurements to ask if: (1) there are significant main effects of treatment (i.e., visit), sex, and/or age, and (2) if there are significant interaction effects (age:sex, age:visit, sex:visit, age:sex:visit) on plasma cytokine levels in our population. We summarize our results below (Figure 3, Supplmental Table S2).

### Pro-inflammatory cytokines

We found a small, but highly significant, overall effect of visit on TNF-*α* levels (*p* = 1.282 × 10^−7^), where treatment (V2) was associated with reduced levels. We also observed a significant overall effect of age (*p* = 1.200 × 10^−7^), where infants had higher overall levels compared with adults, and a marginally significant effect of sex (4.569 × 10^−2^)—males had higher average levels of TNF-*α* in both age groups and time-points. We observed a significant sex-specific effect on IFN-*γ* levels (*p* = 2.048 × 10^−2^), and an age:sex:visit interaction effect (*p* = 3.85 × 10^−3^). We found that IL-6 decreased significantly after treatment (*p* = 1.907 × 10^−2^). Even though we did not detect significant sex-based effects on IL-6, the discordant response we observed between infant and adult samples in males contrasted with the similar response we observed in both age groups in females. We observed a significant overall treatment effect on IL-12(p70) levels (*p* = 3.483 × 10^−6^), where post treatment levels were higher than during acute infection, and a nearly significant sex effect (*p* = 1.291 × 10^−2^) where males had slightly higher values at both time points and in both age groups. We observed no overall effect of age on IL-12(p40) levels, however we found that in males, there appeared to be a treatment effect in adults only, with higher IL-12(p40) levels after treatment, and in females, there appeared to be a treatment effect in infants only, with higher IL-12(p40) levels after treatment. This manifested as a marginal age:sex:treat effect (*p* = 3.475 × 10^−2^).

Observed levels of IL-1,*β* were often below detectable limit, and the levels of sCD40L were often above the detectable range, making their quantification highly uncertain, and leading us to exclude those cytokine measurements from our analysis.

### Anti-inflammatory cytokine and growth factor

We observed a significant effect of visit (treatment) on plasma levels of IL-10 (*p* = 2.566 × 10^−15^), where post treatment levels were substantially lower than during acute infection. We also observed a significant effect of age on IL-10, where infants had significantly higher levels than adults at both time points (*p* = 3.305 × 10^−7^). We observed a small but significant effect of treatment on levels of GM-CSF in the plasma (*p* = 1.151 × 10^−3^), where post-treatment individuals had slightly elevated GM-CSF, regardless of age group. Males trended toward higher mean values across the two time points and ages.

### Treatment failure

Although we did not expect *a priori* for there to be antimalarial treatment failures in our cohort, we found that five individuals remained parasitaemic even after treatment, likely indicating treatment failure and/or reinfection by V2 (Supplemental Figure S3). Among the five, parasite levels were reduced by only ~5% in a single female infant, and by >97% in the remaining 4 individuals. All five individuals had lower plasma IL-10 and TNF-*α* on V2 compared with V1, similar to the general effect across all study participants. However, notable among most of these subjects is the substantial decrease in IL-6 to very low levels on V2.

### Plasma cytokine ratios

The ratios of distinct plasma analytes, many of which simultaneously compete to modify the plasma immune milieu, may more precisely characterize the immune landscape at different levels of treatment, age, or sex. We examined the plasma cytokines, TNF-*α*, IFN-*_γ_*, IL-6, IL-12(p70), IL-10, and GM-CSF, consisting of 15 pairwise analyte combinations, and analyzed effects of age, sex, and visit on their proportions. We found that there were significant overall effects of age in 10 of 15 of the proportions examined. In contrast to effects on individual analyte levels, we observed no overall sex-specific effects on analyte ratios. We found significant overall effects of treatment (visit) on 13 of 15 analyte proportions, and significant age effects on treatment response for five of 15 proportions, with the most significant effects observed on IL-6 / IL-12(p70) treatment response (*p* = 1.385 × 10^−4^) and IL-6 / GM-CSF treatment response (*p* = 8.994 × 10^−4^), where age reversed the direction of the treatment response in both cases (Figure 4, Supplemental Table S3). The most significant sex-dependent age effects on treatment response that we observed were for IFN-*_γ_* / IL-12(p70) (*p =* 8.849 × 10 ^−4^) and IFN-*_γ_* / GM-CSF *(p* = 9.116 × 10^−4^).

**Figure 4:**
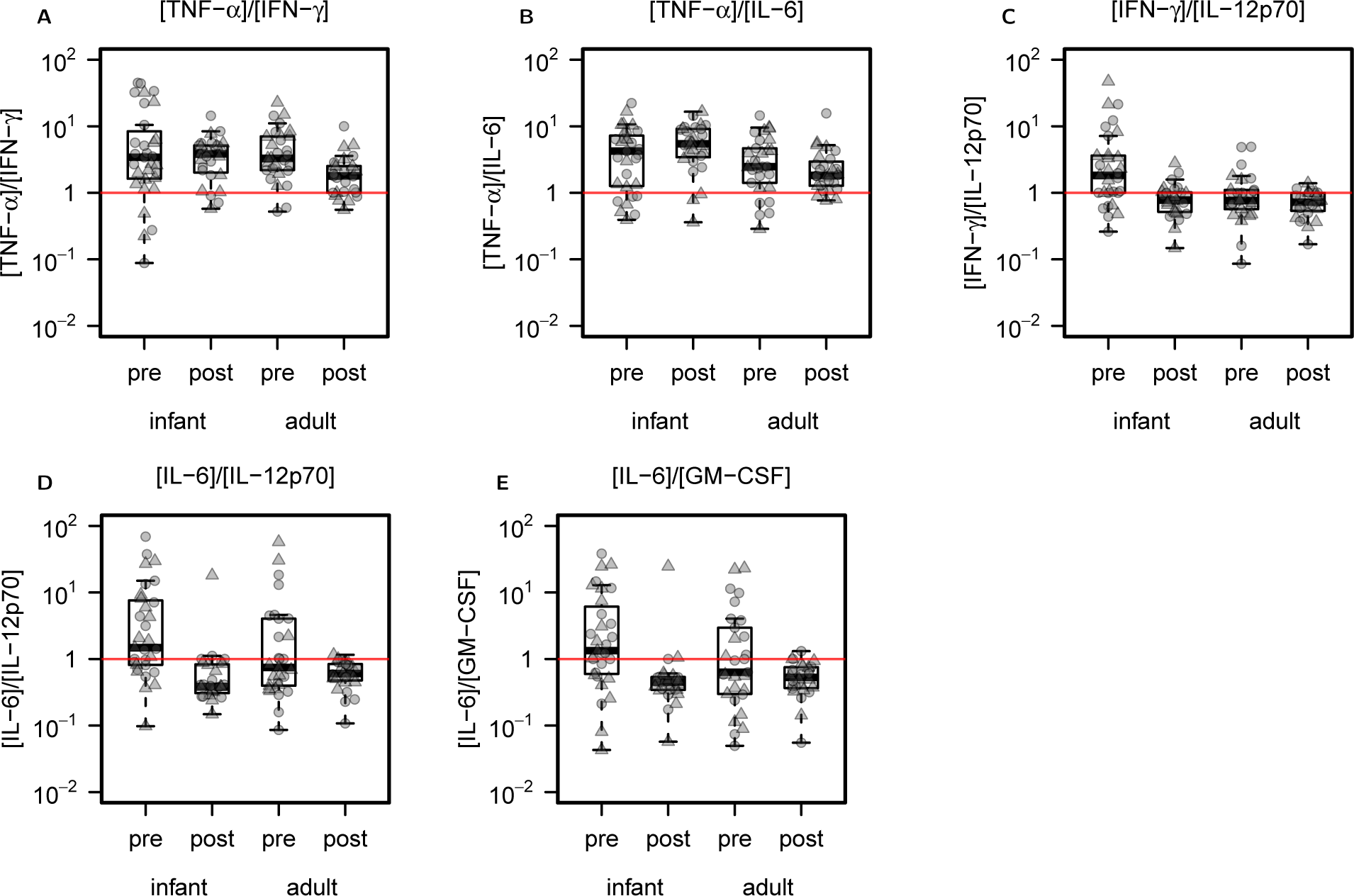
Treatment response of blood analyte ratios is modified or reversed in infants compared with adults. Analyte ratios for which a significant treatment response interaction with age was discovered (5 of 15 tested) are presented as proportions, for: (A) TNF-*α*/IFN-*_γ_*, (B) TNF-*α*/IL-6, (C) IFN-*_γ_*/IL-12(p70), (D) IL-6/IL-12(p70), and (E) IL-6/GM-CSF. The horizontal line demarks a ratio of 1:1.

### Monocyte and Dendritic Cell Composition

Functional differences in immune responses and inflammatory signaling between individuals may be mediated by differences in the overall composition of monocytes and monocyte-derived cellular populations circulating in the blood. We did not observe any significant difference in percentages of CD33^+^ cells based on age, sex, or visit/treatment, however we observed an overall trend for higher percentages in the second visit than during the first, and for higher levels in adults than infants (Figure 5A). We found that the proportion of myeloid dendritic cells (mDCs) among all PMBCs, while very small (often < 0.1%), was significantly higher post-treatment than during acute infection in all groups (*p* = 6.032 × 10^−8^). In addition, nearly significant effects were observed for age (p = 4.665 × 10^−2^) and an age:visit interactions (*p* = 4.282 × 10^−2^), mostly due to lower mDC levels in infants vs. adults during the acute visit (similar levels of post-treatment mDCs) (Figure 5B).

**Figure 5:**
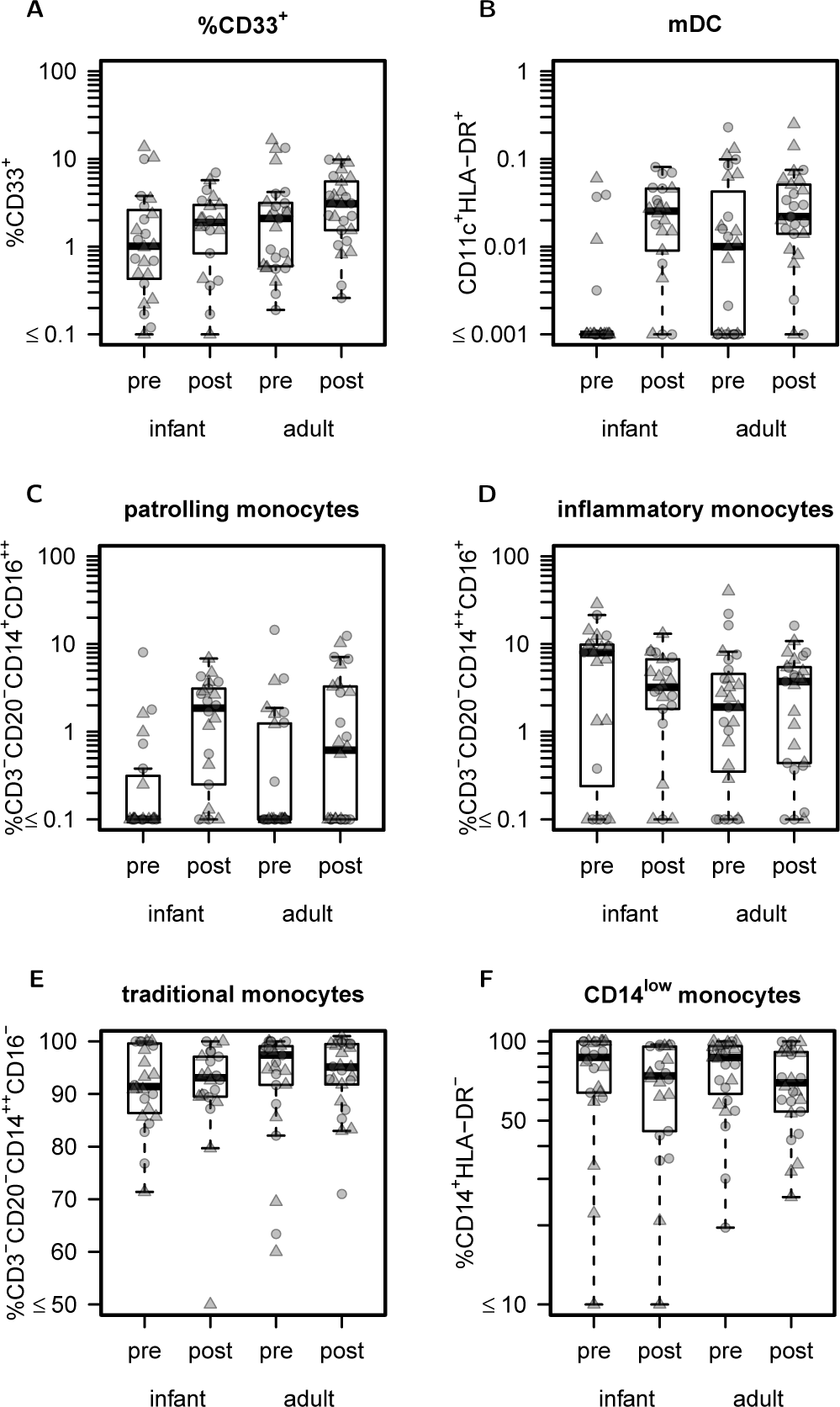
Differences in the composition of myeloid DCs, patrolling monocytes, and CD14^low^ monocytes based on age and/or visit.. The percent composition of (A) CD33^+^ cells, (B) mDCs among all viable PBMCs is shown. The percent composition of (C) patrolling, (D) inflammatory, and (E) traditional monocyte subsets, as a fraction of all monocytes, as well as (F) the percent of CD14^low^ monocytes, as a percentage of all CD16^—^ monocytes, are shown. Percentages are stratified by age group and visit.

Traditional, classical and patrolling monocytes serve different roles in pathogen surveillance, effector functions, and disease pathogenesis [42]. We observed a significant treatment effect on patrolling monocytes (*p* = 1.168^−5^), where levels increased significantly post-treatment in both infant and adult populations (Figure 5C). Although not significant, it appeared that age changed the direction of the treatment response for both inflammatory (Figure 5D) and traditional monocytes (Figure 5E). We observed a significant effect of treatment on the frequency of CD14^low^ monocytes (*p* = 1.648 × 10^−2^) as a percentage of the total CD16 monocytes (Figure 5F).

A summary of p-values for age, sex, visit, and interaction effects for all analyte, analyte ratio, and cellular p-values is included in Figure 6.

**Figure 6:**
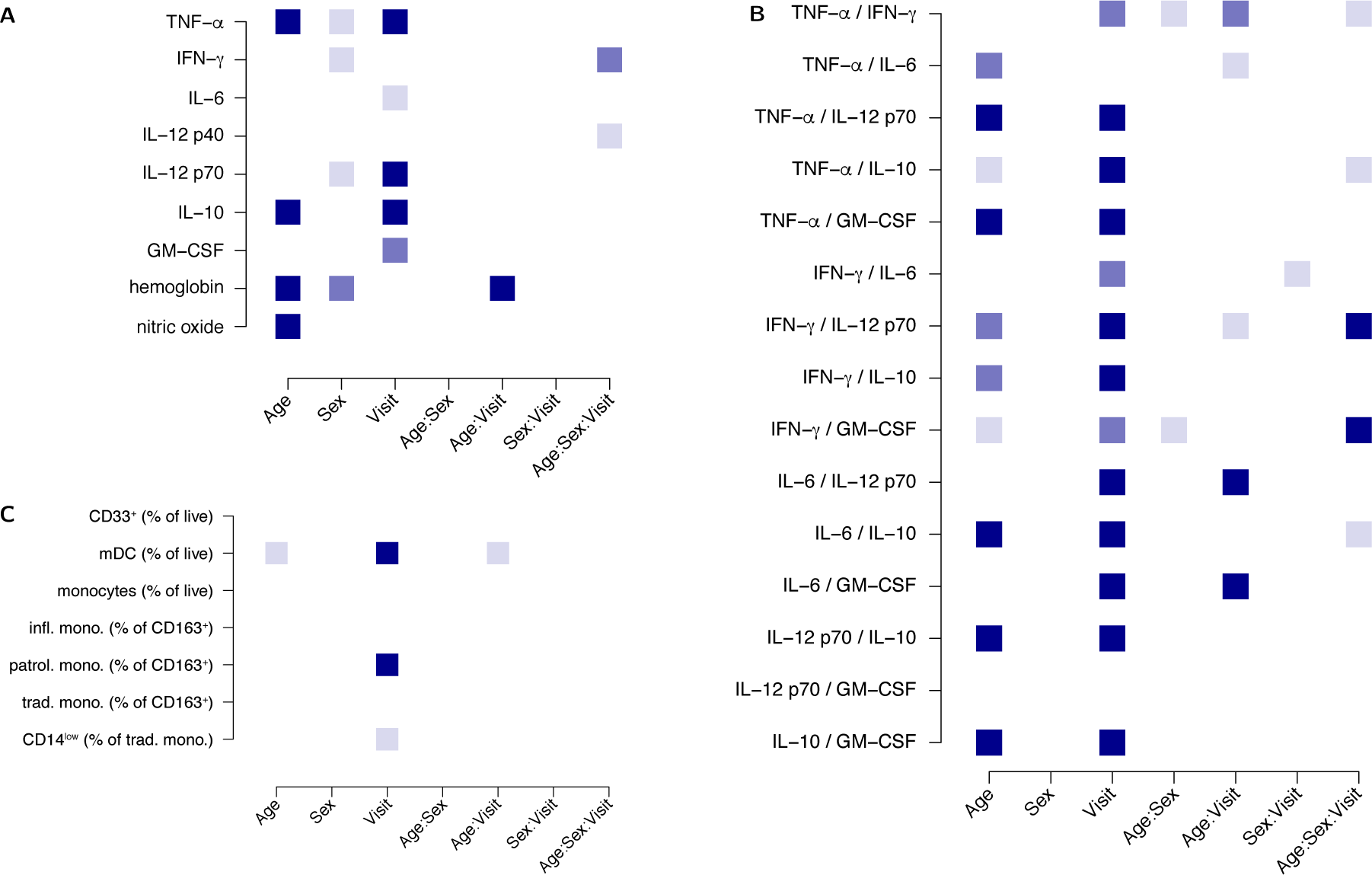
Significant factors and interactions on blood analytes, analyte ratios, and cellular phenotypes identified in this study. Nominal p-values for factors identified by nonparametric analysis of blood analytes (A), analyte proportions (B), and cellular data (C) are indicated by color (light blue: *p <* 0.05; medium blue: *p <* 0.01; dark blue: *p* < 0.001).

### Within-group age-dependent effects on analyte levels

Additional blood analyte heterogeneity within-group, adult or infant, may be caused by age-dependent effects that are not captured by the binary coding of age used in our main analysis. To identify continuous rather than categorical age effects, we used a linear model, fitting age (in years for adults, or fraction of years for infants) and age-by-sex effects for adults and infants separately, at each treatment timepoint, and fitting the same effects for the log_2_-fold change between acute and post-treatment visits. Although we found no significant effects on the treatment response (log_2_-fold change), we identified significant within-group age effects at both visit 1 and visit 2.

At visit 1, we found significant within-group age effects on infant TNF-*α* (ANOVA-like *p* = 0.008, decrease with age, appears to be driven by females) (Figure 7A), and on adult GM-CSF (*p* = 0.032, increase with age), adult IL-12(p70) (*p =* 0.0475, slight decrease with age, not shown), and adult *Pfs16* (*p =* 0.00976, decrease with age), including substantial effects of age (*p* = 0.0032) and age-by-sex interaction (*p* = 0.0027) (Figure 7B).

**Figure 7:**
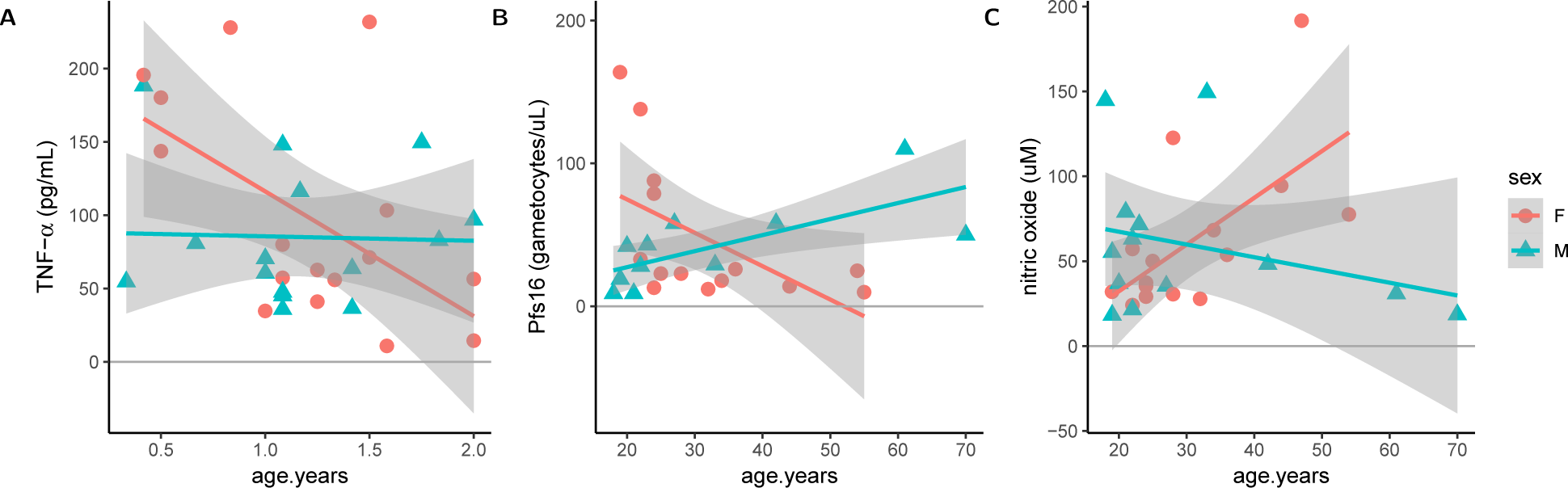
Continuous age is associated with differences in host and parasite factors in both infant and adult populations. Continuous age effects and age-by-sex interaction effects are shown for (A) TNF-*α* at V1 for infants, (B) *Pfs16* levels at V1 for adults, and (C) NO levels at V2 for adults. Age is presented in years.

At visit 2, we found significant within-group age effects on adult nitric oxide (p = 0.014, increase with age), including substantial effects of age (*p* = 0.017) and age-by-sex interaction (*p* = 0.0093) (Figure 7C).

## Discussion

In order to resolve the ongoing problem of malaria transmission in endemic regions, substantial improvements in the quality and/or coverage of preventive measures and treatments are critical. Variation in the host immunological response to infection and treatment may underlie variability in treatment efficacy and clinical outcomes, and infant and immune-compromised populations are especially at risk for adverse outcomes even when adequate antimalarials are available. In this study we evaluated age-related differences in the antimalarial treatment response in adults and infants acutely infected with *Plasmodium falciparum*, the predominant malaria parasite in southern Africa. We provide evidence for substantial widespread differences in immune regulatory factors and cellular effectors between adult and infant populations which are infected with *P. falciparum* and subsequently treated, suggesting that age-related factors may interfere with *both* host-intrinsic anti-parasite immunity as well as the host response to anti-parasite chemotherapy. This two-fold involvement of age effects presents an obstacle to potential vaccine and drug-mediated interventions to eliminate the transmission of malaria, especially in malaria endemic regions of Africa.

Immunological ontogeny and maturation is a process that is highly developmentally and environmentally regulated, and immune features, both at baseline and in response to stimuli, are subject to non-linear trajectories over a typical human lifespan. One of the major factors driving age-dependent differences in the treatment response to antimalarial drugs is immunity of the human host [43]. Thus we focused our efforts on understanding the immune response, based on measures collected in peripheral blood. In our study of age-specific effects on infection and treatment responses of infants and adults, we found substantial effects of age on blood-stage parasitemia, gametocytemia, and a greater risk of recrudescence or reinfection. In addition, blood marker levels were significantly different in infants compared with adults during acute infection, and changes in these levels in response to treatment also differed. When considering co-variation of blood analytes, in the form of cytokines ratios, we found that infant age modifies or reverses the effects of treatment. We observed age-related differences in the treatment response of myeloid DCs. Finally, we observed that within infant or adult age groups, continuous age effects, and age-by-sex effects, contributed to phenotypic differences observed at V1 and V2, sometime transgressing the observed group-wise age effects, shedding a light on the complexity of immune development at long-term and short-term time scales.

In our study, all subjects were infected, based on clinical features and rapid diagnostic test. However, among all infected individuals, we found that infant age was associated with increased numbers of mature gametocytes during acute infection, and decreased overall parasite clearance post-treatment. Infants exhibited significantly higher parasite loads during the first clinical visit, and higher levels of (*Pfs25-* expressing) mature gametocytes, reflecting potential differences in biology, disease presentation and/or healthcare seeking. It has been shown that transplacentally-transferred antibody decreases over time after birth [44]. We found total anti-Plasmodium IgG and IgM levels were detectable at a lower frequency in infants compared with adults, potentially conferring lower or higher levels of protection from pathology in infected individuals. Even so, the majority of infants tested positive, suggesting high rates of prior exposure in infants and/or retention of substantial detectable maternal antibody.

The role of nitric oxide (NO) in malaria is still controversial. Even as higher levels of NO are associated with CM and dyserythropoiesis, some studies have found that NO levels are often higher in asymptomatic children. Our results suggest that NO levels are upregulated in infants compared with adults, however these measures did not change between V1 and V2, and they did not correlate with parasitemia as other studies have found [45]. This may reflect our power to detect such effects given the size of our sample, or differences in regional or environmental factors contributing to NO levels in the blood.

One limitation of our current study is that we lacked clinical outcomes beyond a simple measure of parasitemia, limiting our ability to understand the impact of our blood phenotyping results on adverse outcomes; we excluded individuals with signs of SMA or CM, so the clinical variation in this study is by design very low. Even so, we were able to identify specific signatures of infant age that are associated with changes in the host antimalarial treatment response, flagging factors for future follow-up. Identifying categorical or quantitative clinical measures that differentiate successful treatment from unsuccessful treatment in different age groups could lead to more detailed recommendations for improving treatment protocols in vulnerable host populations.

Some important questions remain. Going forward, it will be important to understand not only the range of treatment responses between age groups, but the important and remediable age-associated factors that lead to recrudescence and/or rapid reinfection. Infants may have altered pharmokinetics, tend to vomit doses of medicine, and/or have differential adherence to treatment compared with adults. Genotyping or sequencing of parasites in future studies will enable us to distinguish treatment failures from new infections. Carefully designed studies that take these factors into account more closely will help us to reduce risk of poor treatment outcomes in pediatric populations more effectively.

Information about prior clinical exposure will be important to consider, since even in areas of high malaria transmission, substantial heterogeneity of exposure is possible, regardless of age group. Indicators of current, or recent, parasite infection may differ substantially, due to the differences in sensitivity and specificity of the assays conducted. A recent analysis of malaria rapid diagnostic tests suggests around 95% specificity and 95% sensitivity of these assays [46]. However, as the antigens detected in these tests are often present even after effective treatment, they are not useful for determining frequency of treatment failures, and parasite microscopy methods are preferred. We used thick film microscopy at V1 and V2 to determine levels of parasite clearance, and in future work with larger sample sizes, more highly quantitative measurements can be taken to better characterize treatment failure rates. Several of our subjects had levels of parasites at V1 that were near or above the threshold considered *hyperparasitemic* (>400,000 parasites/ul). At this level, uncomplicated hyperparasitemia results in higher treatment failure rates, and indeed we observed treatment failures in those with some of the highest parasite levels at visit 1. Hospitalization may sometimes be required for individuals found to be parasitemic at this level, regardless of the level of other symptoms. Although we do not know whether the 5 individuals with detectable parasitemia on V2 were treatment failures or recurrences, evidence from this study can be used to inform future studies to test whether the associated cytokine signals are important risk factors for treatment failure and/or susceptibility to re-infection. Experimental animal models of malaria infection and antimalarial treatment may provide us with the ability to determine causality of age effects on differences in treatment outcomes, and specifically understand the pathways to recrudescence.

In our data, we found that at V1, twelve of the 63 individuals had family members with recent or current cases malaria (data not shown). We did not focus on identifying potential parasite transmission pathways, or family or location-specific risk factors that may also have played a role in the response to infection and treatment in this study, although we know they likely contributed to variation in disease outcomes. In addition, co-morbidities/co-infections were not reported, but may have noteworthy effects on post-infection interventions. Thus, genetic, environmental, and family-relatedness measures, as well as additional health record information, can be informative and influential in determining treatment outcomes, and is recommended to collect these data if at all feasible.

Age effects are important for pediatric chemotherapies more broadly, making infant-specific therapies an important focus for combating global pathogens and improving global health. Treatment failures in infants are may complicate a number of infections, such as pneumococcal infection, HIV infection, and others. Thus, in this work, we have helped to identify the role of both categorical age and continuous age on immune-related differences in the response to drug treatment after acute infection, with implications for how we understand the dynamics of immune development and effects of immune exposure to a broad range of pathogens across the lifespan.

It is clear that age plays a role in the eventual outcomes of complicated and uncomplicated malaria treatment. Prior studies have identified age-dependent treatment effects on infection recurrence [47] and treatment failure for a number of antimalarial drugs [48, 49, 50, 51, 52, 43]. Aside from age, there are a number of factors that may contribute to variability in antimalarial treatment. Additional factors include level of regional malaria endemicity or prevalence, nutrition, immune status, prior infection with malaria, chemotherapeutic choice, and other co-infections or co-morbidities. The relative importance of these factors in driving the heterogeneity in antimalarial treatment responses is still to be understood.

## Conclusion

In summary, our data shows that there are signatures from peripheral blood biomarkers that may indicate or mediate immune response differences infants and adults in a malaria endemic region. These differences in important inflammatory cytokines and cell populations may drive the clinical differences observed in disease risk between infants and adults, and furthermore sex effects may play a modifying role. Finally, the lack of efficacy of antimalarial therapy in some individuals, caused by incomplete clearance or repeat infection, may be a function of cytokine dysregulation in the host response, and identification of the regulatory pathways that are altered will be critical to improving chemotherapy outcomes in infants.

## Declarations

### Ethics Approval and Consent to Participate

The study was approved by the UNC-Chapel Hill Institutional Review Board (# 11-1906) and by the National Health Sciences Research Committee (NHSRC; # 882) that is part of the Ministry of Health in Malawi, in Lilongwe, Malawi. Informed parental consent was obtained. Institutional guidelines at UNC and at the UNC Project Malawi in Lilongwe, Malawi, strictly adhere to the World Medical Association’s Declaration of Helsinki.

### Consent for publication

Not Applicable.

### Availability of data and material

The data file and analysis scripts we be made available in the **malariaInfantStudy** repository on GitHub at https://github.com/mauriziopaul/malariaInfantStudyupon publication.

### Author’s contributions

KDP conceived and designed the experiments. PLM, HF, and GT performed the experiments. MH oversaw the clinical studies in Malawi. PLM, HF, and KDP analyzed the data and wrote the paper, with feedback from all authors.

#### Acknowledgements

We acknowledge the assistance of Dr. Hyung-Suk Kim, from UNC Department of Pathology and Laboratory Medicine (retired), with the gametocytemia PCR methods.

## Supplemental Materials for Malaria Infant Study

### Supplemental Methods

#### Supplemental Statistical Methods

##### Parametric analysis of analytes

In order to properly transform our data prior to using a linear mixed model, we first handled heteroskedasticity of the residuals using a power transform based on a Box-Cox analysis. We start by fitting our data using a linear model:

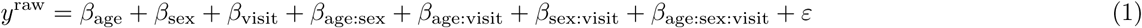

where the values for *ε* are i.i.d. normal. We then use this fit to determine the value of *λ* with the maximum log likelihood, where *λ* represents the exponent and divisor in the follwing data transform:

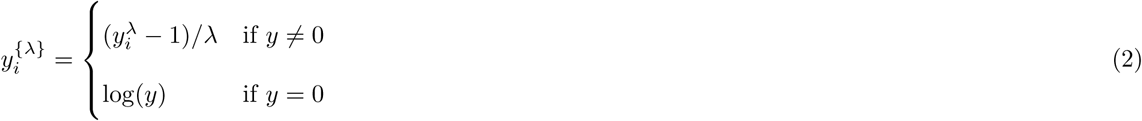

where log is the natural log, according to [53], implemented in the `R` package `MASS` [31]. After finding approximate optimal values of lambda, the transforms used for each phenotypes are as follows, where fractions represent the values of lambda used, “none” represents no transformation, and “log” is the natural log transformation:

| analyte: | GM-CSF | IFN-γ | IL-10 | IL-12(p40) | IL-12(p70) | IL-6 | TNF-α | hemoglobin | nitric oxide |
| --- | --- | --- | --- | --- | --- | --- | --- | --- | --- |
| transform: | -1/3 | -1/2 | -1/3 | -1/3 | [log] | -1/2 | [log] | [none] | [log] |

The quantile-quantile (Q-Q) plots are shown before and after the power transformations are applied in Figure S4. We then applied a linear mixed model to the transformed data, according to the same model in equation (1), but adding individual-level random slopes (for visit).

#### Supplemental Results

##### Results for sCD40L and IL-1ß

Soluble CD40 ligand (sCD40L, also known as sCD154) has been shown to induce DC maturation following *Plasmodium* infection in vitro, which can result in release of TNF-*α* and IL-12(p70) [54]. We observed a marginal visit effect (*p =* 8.783 × 10^−3^), with higher levels post-treatment, as well as a marginal sex-specific effect on sCD40L (*p* = 6.734 × 10^−3^), with higher overall levels in males. However, the distribution of phenotypes was near the upper detection limit for this cytokine, which may have reduced our ability to detect differences based on treatment, age, and sex. We found that IL-1*β* levels in plasma increased significantly after treatment (*p =* 9.162 × 10^−4^), and we found no other significant effects. This cytokine was at the lower limit of detection, which may have also reduced our ability to detect significant effects.

##### Linear mixed model analysis of analytes

We expect subtle differences in our main results based on the model that is selected. To determine whether our findings are sensitive to model type and specification, we applied a linear mixed model in addition to the nonparametric model reported in the main results. We found that the majority (9 of 16) of the significant effects identified by our nonparametric model, presented in Figure 6, were also found using a LMM, as described above (Supplemental Table S4). Using ANOVA on the LMM (using the package `lmerTest`), we uncovered an additional 3 significant effects for IL-6 (sex, *p* = 0.041) and IL-10 (age, *p* = 0.0096; age-by-sex, *p* = 0.020) [55].

**Figure S1:**
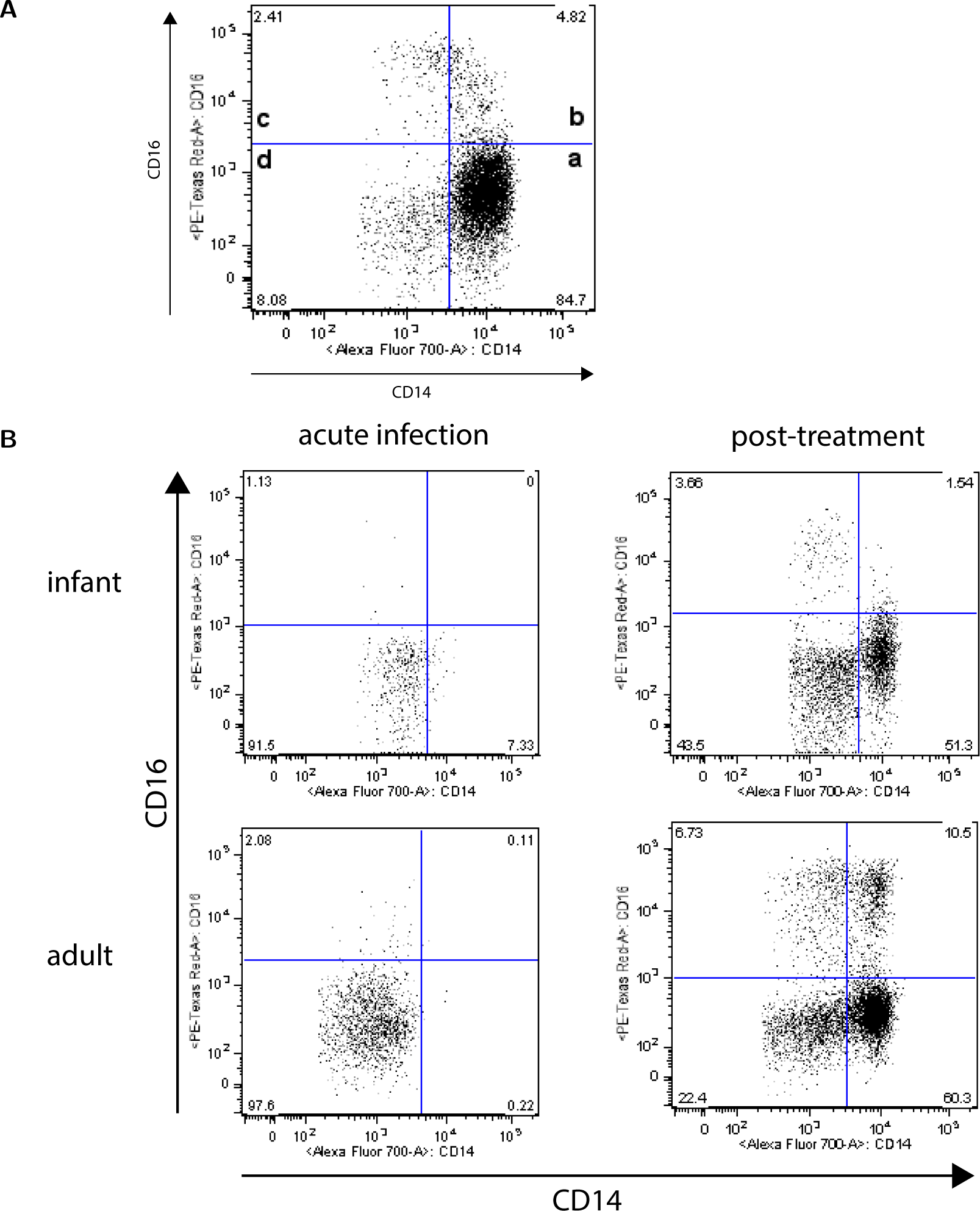
Manual gating strategy for quantifying monocyte subsets. (A) Monocyte cell counts and percentages are derived for CD14^++^CD16^—^ traditional monocytes, as designated by quadrant (a), CD14^++^CD16^+^ inflammatory monocytes, as designated in quadrant (b), and CD14^+^, CD16^++^ patrolling monocytes, as designated in quadrant (c), shown here for an example healthy, uninfected adult. A population of monocytes with intermediate CD14 brightness is designated in quadrant (d). (B) Example of flow cytometric phenotypic characterization of monocyte subsets in a *Plasmodium-infected* infants and adults is shown before and after treatment.

**Figure S2:**
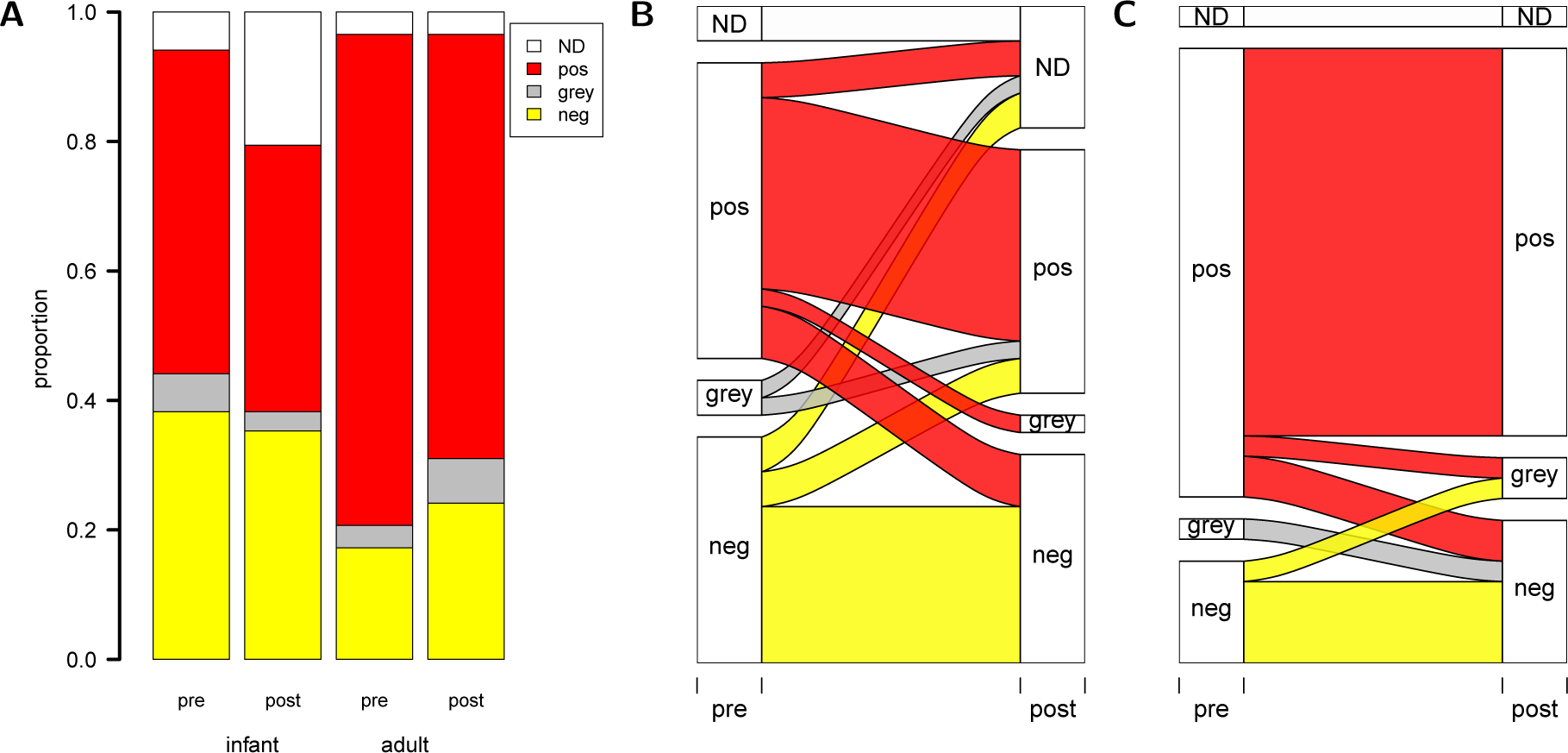
Plasmodium-specific antibody levels higher in adults compared with infants regardless of visit. (A) The test result proportions for overall circulating antimalarial IgG and IgM, measured by ELISA test of blood samples, are shown. The change in test result status between visit 1 and 2 is shown for (B) infants and (C) adults. Colors indicate positive (red), uncertain (grey), and negative (yellow) test results.

**Figure S3:**
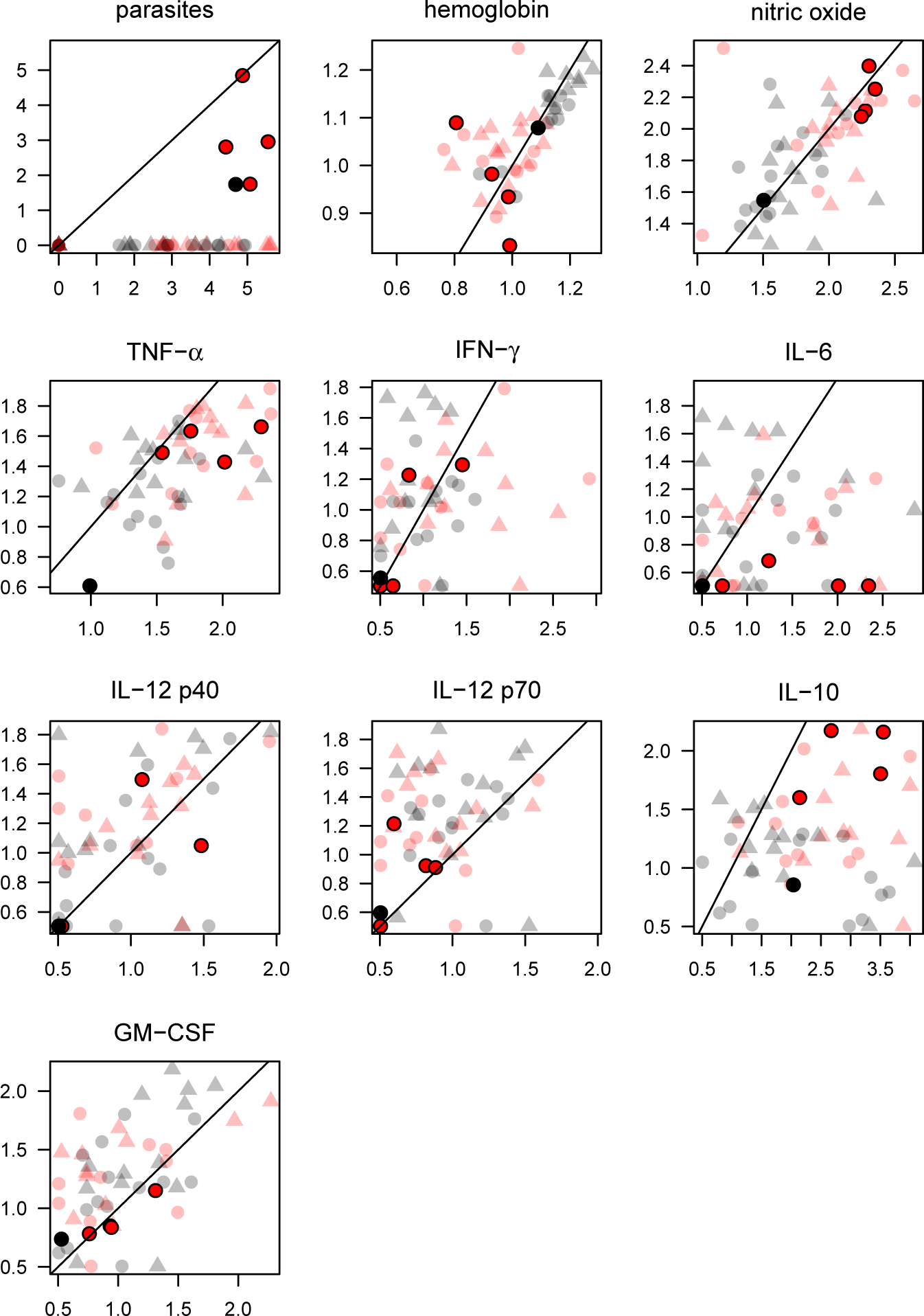
Correspondence between acute and post-treatment phenotypes. The log (base=10) values of phenotypes from Figure 2A and Figure 3 are presented, with values at Visit 1 (acute) provided along the x-axis, and values for Visit 2 (post-treatment) along the y-axis. The black line in each figure indicates *y = x*. Circles indicate females and triangles indicate males. Pink or red indicates infants and grey or black indicates adults. The five darker colored dots in each plot indicate phenotypes for individuals with apparent treatment failure.

**Figure S4:**
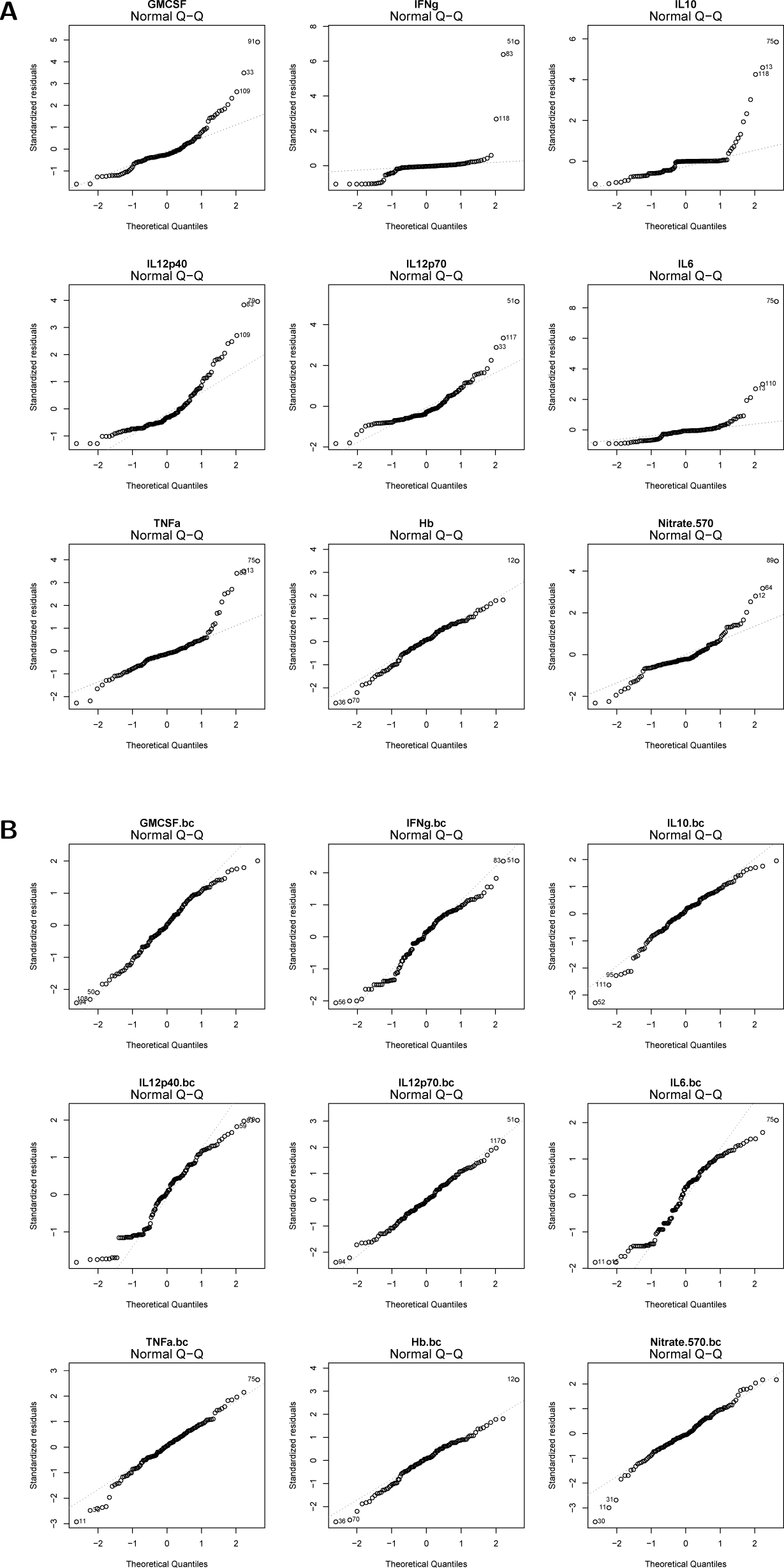
QQ-plots of analyte phenotypes (A) before and (B) after Box-Cox-assisted power transformation.

**Table S1:** Results from antimalarial antibody detection. Values indicate the proportion of samples within each age-visit group that received a given result for total Ig (IgG and IgM), where: "grey" indicates that the sample result was indeterminate and "N/A" indicates that the sample or result is missing.

|  | Result | adult-visit-1 | adult-visit-2 | infant-visit-1 | infant-visit-2 |
| --- | --- | --- | --- | --- | --- |
| 1 | positive | 0.793 | 0.655 | 0.500 | 0.441 |
| 2 | grey | 0.035 | 0.069 | 0.059 | 0.029 |
| 3 | negative | 0.172 | 0.241 | 0.382 | 0.382 |
| 4 | N/A | 0.000 | 0.035 | 0.059 | 0.147 |
| 5 | Total | 1.000 | 1.000 | 1.000 | 1.000 |

**Table S2:** Table of *p*-values for main effects and interaction effects on blood analytes and cellular phenotypes, analyzed using a nonparametric longitudinal model.

| Analyte | Age | Sex | Visit | Age:Sex | Age:Visit | Sex:Visit | Age:Sex:Visit |
| --- | --- | --- | --- | --- | --- | --- | --- |
| TNF-α | 1.200e-07 | 4.569e-02 | 1.282e-07 | 3.633e-01 | 4.842e-01 | 5.443e-01 | 7.726e-01 |
| IFN-γ | 1.391e-01 | 2.048e-02 | 8.164e-01 | 1.844e-01 | 2.069e-01 | 8.839e-01 | 3.851e-03 |
| IL-6 | 4.632e-01 | 6.416e-02 | 1.907e-02 | 6.059e-01 | 5.987e-02 | 6.954e-02 | 4.721e-01 |
| IL12(p40) | 3.549e-01 | 2.410e-01 | 8.155e-02 | 9.803e-01 | 6.300e-01 | 8.602e-01 | 3.475e-02 |
| IL12(p70) | 2.641e-01 | 1.291e-02 | 3.483e-06 | 9.633e-01 | 4.315e-01 | 3.689e-01 | 9.690e-01 |
| IL-10 | 3.305e-07 | 3.055e-01 | 2.566e-15 | 5.364e-01 | 9.228e-01 | 6.009e-01 | 7.045e-02 |
| GM-CSF | 4.867e-01 | 6.934e-02 | 1.151e-03 | 3.555e-01 | 4.518e-01 | 4.510e-01 | 2.546e-01 |
| hemoglobin | 3.860e-15 | 5.592e-03 | 2.845e-01 | 7.950e-02 | 3.134e-04 | 8.303e-01 | 8.445e-01 |
| nitric oxide | 1.191e-10 | 6.485e-01 | 8.767e-01 | 5.313e-01 | 6.629e-01 | 1.525e-01 | 5.243e-01 |
| CD33<sup>+</sup>, % of live | 1.234e-01 | 3.144e-01 | 6.351e-02 | 8.146e-01 | 5.770e-01 | 3.976e-01 | 5.409e-01 |
| mDC, % of live | 4.666e-02 | 9.955e-01 | 6.032e-08 | 2.024e-01 | 4.282e-02 | 7.180e-01 | 7.882e-01 |
| monocytes, % of live | 1.903e-01 | 3.264e-01 | 1.303e-01 | 9.151e-01 | 4.617e-01 | 3.037e-01 | 5.550e-01 |
| infl. monocytes, % of CD163<sup>+</sup> | 1.269e-01 | 3.639e-01 | 7.735e-01 | 9.002e-01 | 1.780e-01 | 5.000e-01 | 5.468e-01 |
| patr. monocytes, % of CD163<sup>+</sup> | 7.971e-01 | 4.551e-01 | 1.168e-05 | 3.814e-01 | 1.104e-01 | 2.660e-01 | 9.464e-01 |
| trad. monocytes, % of CD163<sup>+</sup> | 1.072e-01 | 1.738e-01 | 7.886e-01 | 9.705e-01 | 8.510e-01 | 6.950e-01 | 9.854e-01 |
| CD14<sup>low</sup>, % of traditional | 4.337e-01 | 9.392e-01 | 1.648e-02 | 2.313e-01 | 4.256e-01 | 3.040e-01 | 2.756e-01 |

**Table S3:** Table of *p*-values for main effects and interaction effects on blood analyte ratios.

| Analyte Proportion | Age | Sex | Visit | Age:Sex | Age:Visit | Sex:Visit | Age:Sex:Visit |
| --- | --- | --- | --- | --- | --- | --- | --- |
| TNF-α/IFN-γ | 9.121e-02 | 5.852e-01 | 2.680e-03 | 2.673e-02 | 8.262e-03 | 5.840e-01 | 1.691e-02 |
| TNF-α / IL-6 | 2.857e-03 | 6.443e-01 | 6.791e-01 | 9.455e-02 | 2.693e-02 | 1.370e-01 | 6.000e-01 |
| TNF-α / IL12(p70) | 5.861e-08 | 4.937e-01 | 3.807e-18 | 3.270e-01 | 4.423e-01 | 3.774e-01 | 4.588e-01 |
| TNF-α / IL10 | 2.706e-02 | 6.958e-01 | 4.555e-16 | 8.375e-01 | 2.540e-01 | 9.063e-01 | 4.831e-02 |
| TNF-α / GM-CSF | 1.907e-05 | 4.098e-01 | 8.120e-13 | 8.466e-01 | 5.674e-01 | 9.900e-01 | 2.218e-01 |
| IFN-γ / IL-6 | 3.077e-01 | 9.418e-01 | 1.782e-03 | 9.354e-01 | 3.535e-01 | 1.892e-02 | 3.071e-01 |
| IFN-γ / IL-12(p70) | 1.128e-03 | 7.762e-01 | 5.501e-06 | 1.244e-01 | 2.191e-02 | 4.324e-01 | 8.849e-04 |
| IFN-γ / IL-10 | 1.034e-03 | 6.961e-01 | 2.844e-19 | 6.569e-01 | 6.257e-01 | 8.150e-01 | 5.466e-01 |
| IFN-γ / GM-CSF | 2.533e-02 | 3.089e-01 | 1.093e-03 | 3.406e-02 | 3.338e-01 | 9.142e-01 | 9.116e-04 |
| IL-6 / IL-12(p70) | 2.524e-01 | 8.248e-01 | 1.500e-11 | 6.402e-01 | 1.385e-04 | 3.185e-01 | 1.507e-01 |
| IL-6 / IL-10 | 5.282e-05 | 3.498e-01 | 3.055e-19 | 8.788e-01 | 5.374e-01 | 1.644e-01 | 1.252e-02 |
| IL-6 / GM-CSF | 3.188e-01 | 5.405e-01 | 1.493e-10 | 2.702e-01 | 8.994e-04 | 7.862e-02 | 3.733e-01 |
| IL-12(p70) / IL-10 | 7.763e-06 | 5.190e-01 | 4.754e-22 | 8.536e-01 | 5.150e-01 | 7.358e-01 | 1.019e-01 |
| IL-12(p70) / GM-CSF | 6.258e-01 | 9.159e-01 | 9.159e-02 | 2.446e-01 | 4.148e-01 | 8.555e-01 | 3.113e-01 |
| IL-10 / GM-CSF | 2.796e-05 | 4.987e-01 | 1.242e-22 | 7.694e-01 | 6.191e-01 | 8.830e-01 | 7.485e-02 |

**Table S4:** Table of *p*-values for main effects and interaction effects on blood analytes, analyzed using a linear mixed model. P-values for model intercepts were all highly significant (ranging in scale from 10^−17^ to 10^−31^), and were omitted from this table.

| Analyte | Age | Sex | Visit | Age:Sex | Age:Visit | Sex:Visit | Age:Sex:Visit |
| --- | --- | --- | --- | --- | --- | --- | --- |
| TNF-α | 7.852e-05 | 4.299e-03 | 1.682e-02 | 6.426e-02 | 6.774e-01 | 5.274e-01 | 6.689e-01 |
| IFN-γ | 4.406e-01 | 8.479e-02 | 9.331e-01 | 5.255e-01 | 7.562e-01 | 2.155e-01 | 6.223e-02 |
| IL-6 | 6.177e-01 | 4.127e-02 | 2.161e-01 | 8.035e-01 | 2.432e-01 | 1.480e-01 | 6.531e-01 |
| IL-12(p40) | 1.144e-01 | 1.068e-01 | 9.536e-01 | 2.881e-01 | 2.896e-01 | 2.898e-01 | 1.201e-01 |
| IL-12(p70) | 9.197e-01 | 6.301e-02 | 3.518e-01 | 7.654e-01 | 7.349e-01 | 3.455e-01 | 5.923e-01 |
| IL-10 | 2.306e-05 | 9.613e-03 | 4.598e-07 | 1.986e-02 | 2.178e-01 | 8.230e-02 | 6.527e-02 |
| GM-CSF | 9.948e-01 | 1.352e-01 | 3.964e-01 | 7.037e-01 | 8.984e-01 | 9.816e-01 | 8.491e-01 |
| hemoglobin | 3.866e-02 | 1.711e-02 | 3.956e-01 | 1.505e-01 | 3.248e-02 | 6.111e-01 | 8.220e-01 |
| nitric oxide | 5.558e-04 | 8.707e-01 | 5.654e-01 | 6.656e-01 | 8.911e-01 | 3.711e-01 | 7.026e-01 |

